# OTTER, a new method quantifying absolute amounts of tRNAs

**DOI:** 10.1101/2020.05.18.101501

**Authors:** Akihisa Nagai, Kohei Mori, Yuma Shiomi, Tohru Yoshihisa

## Abstract

To maintain optimal proteome, both codon choice of each mRNA and supply of aminoacyl-tRNAs are two principal factors in translation. Recent reports revealed that tRNAs are more dynamic in amount than we had expected. High-throughput methods such as RNA-Seq or microarray are versatile for comprehensive analyses of changes in individual tRNA amounts, but they suffer from lack of assessment of signal production efficiency of individual tRNA species. Thus, they are not the perfect choice to measure absolute amounts of tRNA. Here, we introduce a novel method for this purpose, oligonucleotide-directed three-primer terminal extension of RNA (OTTER), which employs fluorescence labeling at the 3’-terminus of a specific tRNA by optimized reverse primer extension and an assessment step of each labeling efficiency by Northern blotting. We quantified absolute amounts of 34 individual and 4 pairs of isoacceptor tRNAs out of the total 42 nuclear-encoded isoacceptors in the yeast *Saccharomyces cerevisiae*. We revealed that the amounts of tRNAs are in the range of 0.030–0.73 pmol/μg RNA in the yeast cells logarithmically grown in a rich glucose medium. The tRNA amounts seem to be regulated at the isoacceptor level by a few folds according to physiological growing conditions. Data obtained by OTTER are poorly correlated with those by simple RNA-Seq and only marginally with those by microarray. However, the OTTER data are good agreement with those by 2D-gel analysis of *in vivo* radiolabeled RNA samples. Thus, OTTER is a suitable method for quantifying absolute amounts of tRNAs in the isoacceptor resolution.

## INTRODUCTION

Building of appropriate proteomes under various conditions is one of the essential tasks for living organisms. Although a repertoire of mRNAs is crucial for building proteomes, supply of aminoacyl-tRNAs determined by a tRNA repertoire and replenishment speed of aminoacyl-tRNAs also have pivotal roles. Thus, selective usage of a particular codon among synonymous ones, in other words, selection of a certain isoacceptor tRNA for peptide bond formation, is important (Brar 2016; Shi and Barna 2015). Indeed, recent progresses in translatome analyses, especially those via ribosome profiling (Ingolia et al. 2009), have revealed a global view that codon selection affects both efficiency (Goodarzi et al. 2016; Qian et al. 2012; Shah and Gilchrist 2011) and accuracy (Drummond and Wilke 2008; Krisko et al. 2014) of translation. Realtime monitoring of protein folding of nascent chains also has been demonstrated that protein folding of a polypeptide encoded by the mRNA is strongly affected by codon selection (Brar 2016; Kim et al. 2015; Yu et al. 2015). Furthermore, recent reports showed that codon selection determines stability of mRNAs under certain conditions (Hia et al. 2019; Mishima and Tomari 2016; Presnyak et al. 2015).

Codon optimality has been usually discussed on codon usage of a gene, and it has been thought that preference of a codon in the genome of an organism in question is paralleled to the amount of a cognate tRNA (Hanson and Coller 2018; Rak et al. 2018). In eukaryotes, tRNAs are usually encoded by multiple genes to produce enough amounts of individual tRNA species. Indeed, tRNAs are the most abundant class of RNAs (Kirchner and Ignatova 2015), and it has been believed that amounts of isodecoder tRNAs, a class of tRNAs with the same anticodon sequence, are mostly determined by numbers of corresponding genes on the genome, meaning that the tRNA repertoire in an organism is constant during its life.

However, several recent reports have casted a doubt on this assumption: Over-expression of initiator or elongator tRNA-Met differently affects tRNA repertoires in human cell lines (Pavon-Eternod et al. 2013). Pilpel’s group compared tRNA abundance in B cell lymphoma with that in normal B lymphocytes, and found that significant expression variations (>10-fold) were seen in certain tRNA species (Gingold et al. 2014). Especially, a pair of isoacceptor tRNAs, tRNA-Lys_CUU_ and tRNA-Lys_UUU_, responded oppositely despite the fact that they decode codons for the same amino acid. Reports of similar change of tRNA repertoires along with transformation and development have been accumulating in these years (Goodarzi et al., 2016; Schmitt et al., 2014; Turowski et al., 2016). These facts indicate that the tRNA repertoire is not a fixed parameter for an organism but is variable according to environmental and developmental conditions. To understand effects of global and local translational efficiency on various biological aspects, it is essential to know concentration of individual tRNA species in cytosol (or in genome-containing organelles) and to obtain a total view of a tRNA repertoire in the cell under certain conditions at the certain time point.

First choice for tRNA repertoire analysis is RNA-Seq. However, several steps in RNA-Seq are strongly affected by the nature of each target. One of such steps is reverse transcription (RT): base modifications and RNA secondary structures strongly alter its efficiency. Especially in quantification of tRNAs, their heavy modifications and tight secondary structures hamper efficient or even full-length RT despite the fact that tRNA species are only 75–90 nt in length (Pang et al. 2014; Zheng et al. 2015). Thus, precise estimation of tRNA amounts by simple RNA-Seq is difficult. It is also true to another RT-based quantification, qRT-PCR (Torrent et al. 2018). Rather, the blockades of RT by base modifications can be positively utilized to identify modified bases in RNA species (Arimbasseri et al. 2015; Hauenschild et al. 2015; Wulff et al. 2017). Several methods have been invented to circumvent the problems in RT. One simple way is to utilize only a 3’ part of tRNA sequence for quantification (Chen and Tanaka, 2018). This method can be applied to organisms with simple genomes, such as *Saccharomyces cerevisiae*. The second choice is usage of RTase with high tolerance to nucleotide modification (Nottingham et al. 2016). Usage of de-modification enzymes to reverse the modification blockades is the third way. Especially, demethylases are adopted to remove the methyl moieties on amino groups of nucleobases with certain success (Zheng et al. 2015). A different approach to avoid modification problems is to introduce additional sequencing start sites by randomly digesting tRNAs by alkaline treatment and to enable sequencing of more distal parts to the 3’-end in a tRNA molecule (Arimbasseri et al. 2015; Karaca et al. 2014). Another improvement has been introduced into the adaptor ligation step required for RT. Kirino’s group reported several devices that improve this step, including usage of Y-shape double-stranded primers instead of stem-loop primers (Honda et al. 2015; Shigematsu et al. 2017).

Another way to measure tRNA quantity is to adopt hybridization. Indeed, microarray has been used to measure changes of tRNA amounts in different samples (Dittmar et al. 2004; Dittmar et al. 2006; Gingold et al. 2014). If compared with RNA-Seq-based methods, the resolution of tRNA species is rather low because near cognate tRNA species with a single nucleotide difference in the region used for probe design may cause cross-hybridization, but there is no need of RT. The stint ligation-based method also utilizes two oligo DNAs that hybridize with adjacent regions on a tRNA followed by ligation of these oligo DNAs. Then, the ligated oligo DNAs, hallmark of the tRNAs, are subjected to next-generation sequencing for quantification (Goodarzi et al. 2016). In either case, base modification may still affect hybridization efficiency of the probes so that, in some cases, designing effective oligonucleotide probes for certain tRNAs cannot be simply carried out just from sequence information of the tRNA genes.

More fundamental problem in both RT-based and hybridization-based high-throughput techniques for tRNA quantification is that it is hard to estimate signal generation efficiency of each tRNA species even after adopting a certain improvement. For example, it is difficult to estimate individual efficiency of hybridization between a certain tRNA species and the corresponding probe on the microarray. It is also true in the RT step in RNA-Seq. Therefore, the present data on tRNA quantification can be used to compare expression difference of each tRNA species under different conditions, such as physiological environments, developmental stages, genetical backgrounds, *etc*. On the other hand, they harbor inherent limitation to compare expression difference between different tRNA species in a strict sense. Thus, we still stay one step before knowing an absolute amount of each tRNA species in a biological sample, which is an important parameter to understand biochemistry of translation *in vivo*.

Here, we introduce a novel method, Oligonucleotide-directed Three-prime Terminal Extension of RNA, or OTTER, to measure an absolute amount of a tRNA species at an isoacceptor level. Although this method also relies on hybridization between the 3’-teminal region of a tRNA species and an oligo DNA that act as a DNA template, the reaction extends the tRNA molecule and labels it with one fluorescent nucleotide. Thus, by analyzing urea-PAGE and fluorescence scanning, the absolute tRNA amount of the modified tRNA can be measured while the modification efficiency can be monitored by Northern blotting of the same sample. Using this technique, we measured amounts of 34 individual isoacceptors and 4 pairs of 2 isoacceptors out of 42 isoacceptors in the yeast *Saccharomyces cerevisiae*.

## RESULTS

### Experimental design of a novel method to quantify tRNA

For quantification of an absolute amount of a specific tRNA species, we adopted reverse primer extension to introduce a fluorescent nucleotide directly to the tRNA species. This method consists of two steps: first, we hybridize an oligo DNA specific to an isodecoder/isoacceptor tRNA with a 5’-extension of 5’-dAdTdTdTdT-3’ (Type 1 oligo DNA; Fig. 1A) to generate a 5’-overhang. As shown in Fig. S1, the 3’-terminal region of yeast tRNAs has enough diversity to design isoacceptor specific oligo DNAs for almost all of the tRNAs. One exception is tRNA-Ser_CGA_ and tRNA-Ser_UGA_, whose last 33 nt sequences, including CCA, are the same. When a certain isoacceptor tRNA consists of multiple isodecoders with the different 3’-terminal sequences, we measured the total amount of the isoacceptor tRNA because there are several pairs of isodecoders for an isoacceptor whose 3’-terminal sequences are identical. In the second step, this hybrid is subjected to DNA polymerization reaction by Klenow enzyme without the 3’-5’ activity with the tRNA as an RNA primer and the oligo DNA as a template. Because the reaction mixture contains only dATP and tetramethylrhodamine-dUTP (TMR-dUTP), the polymerization reaction yields a 5-nt-extended tRNA derivative with the 3’ terminal TMR-dU. The fluorescent tRNA derivative is detected and quantified by fluoro-scanning with a TMR-labeled synthetic oligo DNA as a standard (Fig. 1B), so that we can obtain an absolute amount of the tRNA in the sample. Since the labeled tRNA was elongated by 5 nt, the labeled tRNA and the original tRNA are easily separated by urea-PAGE, and labeling efficiency of each reaction is determined by Northern blotting of the target tRNA (Fig. 1C). Northern blotting is also used to evaluate cross-reactions of a template oligo DNA to non-cognate tRNA species with homologous 3’-terminal sequences (see below). By using these values for compensation, this procedure can quantify an absolute amount of an isoacceptor tRNA, of which accurate estimation is difficult in microarray and RNA-Seq analyses.

**Figure1.**
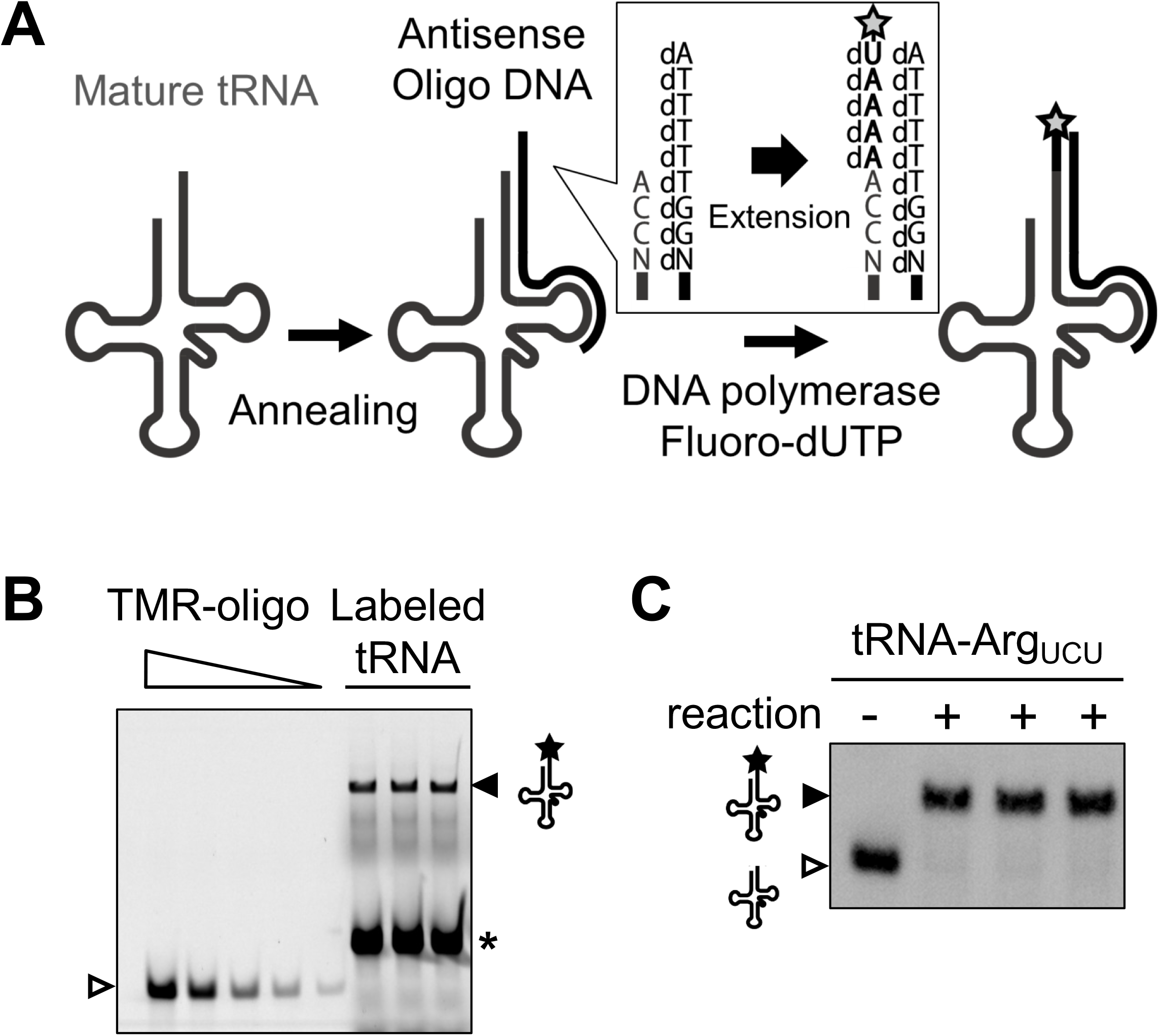
tRNA quantification using OTTER. (*A*) Schematic drawing of tRNA fluorescence labeling reaction in OTTER. A target tRNA is first hybridized with a specifically-designed antisense oligo DNA with the 5’-extension of dAdTdTdTdT (Type 1 oligo DNA). The overhang in the tRNA/DNA hybrid was filled by Klenow fragment (3’-5’ exo^−^) with dATP and fluorescence-labeled dUTP (such as TMR-dUTP; marked by star) as substrates. (*B*) An example for tRNA-Arg_UCU_ quantification. Typical reactions of OTTER for tRNA-Arg_UCU_ were analyzed by urea-PAGE and scanned with a fluorescence scanner. The fluorescence-labeled tRNA-Arg_UCU_ (closed triangle) as well as the fluorescence-labeled template oligo DNA by a weak reverse transcription activity of the Klenow fragment (asterisk) were detected. Since the 3’ end of the oligo DNA falls on the TΨC region rather conserved even among different tRNA species, unrelated tRNAs also acted as templates to produce the strong signal. Three replicates of the reaction were analyzed. The amounts of the standard TMR-oligo DNA on the gel (open triangle) were 0.500, 0.250, 0.100, 0.050, and 0.020 pmol/lane. (*C*) The three OTTER reaction products for tRNA-Arg_UCU_ shown in (*B*) (“+” lanes) were subjected to Northern blotting with a reaction without the template oligo DNA (“−” lane).

Since labeling efficiency of individual tRNAs under the initial assay conditions with Type 1 oligo DNA templates varied significantly among tRNA species (Fig. 2A), we needed to refine assay conditions to each tRNA isoacceptor. During establishment of assay conditions, we mainly encountered three kinds of problems in certain tRNAs: the first problem is incomplete extension where a part of tRNAs ceased to elongate without receiving the final TMR-dU (Fig. 2A, tRNA-Arg_ACG_, asterisk). The second one is low extension efficiency where a considerable portion of certain tRNAs did not receive any extension (Fig. 2A, tRNA-Lys_CUU_, open triangle in the “+” lane). The third is cross-hybridization of certain templates to near-cognate isoacceptor tRNAs (Fig. 2D). The first problem was overcome by redesigning the template oligo DNA to include a dA nucleotide accepting TMR-dU in the middle part but not at the 5’-end of the template (Type 2 oligo DNA; Fig. 2B). Indeed, there was a report that DNA polymerase I interacts with a template strand beyond the site of synthesis (Turner et al. 2003), which may affect efficiency of incorporation of unnatural deoxynucleotides. The second problem seemed to be caused by inefficient hybridization between the target tRNA and the corresponding template oligo DNA coming from structural rigidity of the tRNA. To deal with this problem, we employed an “unfolder” oligo DNA that hybridizes with the anticodon stem-loop and variable loop regions of the target tRNA, and destabilizes its overall structure (Buvoli et al. 2000). As shown in Fig. 2C, introduction of the corresponding unfolder improved extension efficiency in the example of tRNA-Lys_CUU_. For the third problem, extent of cross-reactions to near-cognate tRNAs was monitored by Northern blotting as described previously, and the results were used to optimize annealing temperature to reduce cross-hybridization (Fig. 2D). In the case of tRNA-Val_CAC_, elevating annealing temperature from 30°C to 52°C significantly suppressed the cross-reaction of the oligo DNA to tRNA-Val_UAC_. However, the suppression was not so complete that individual amounts these isoacceptors were calculated from fluorescence signals and labeling efficiency of the cognate and near-cognate isoacceptors (see Materials and Methods in detail). On the other hand, we could not resolve quantitative data of three pairs of isoacceptors, tRNA-Gln_CUG_/tRNA-Gln_UUG_, tRNA-Glu_CUC_/tRNA-Glu_UUC_, and tRNA-Pro_AGG_/tRNA-Pro_UGG_, with satisfactory quantitativeness. As described above, tRNA-Ser_CGA_ and tRNA-Ser_UAG_ cannot be resolved from their homology by our method. Thus, we had no choice to measure the tRNA amounts of these four pairs as the sum of the two isoacceptors. Finally, we could construct a set of measurement conditions for 34 individual and 4 pairs of isoacceptor tRNAs out of the total 42 isoacceptors of *S. cerevisiae* (Table S2). We named this method as <u>O</u>ligonucleotide-directed <u>T</u>hree prime <u>T</u>erminal <u>E</u>xtension of <u>R</u>NA, OTTER in short.

**Figure 2.**
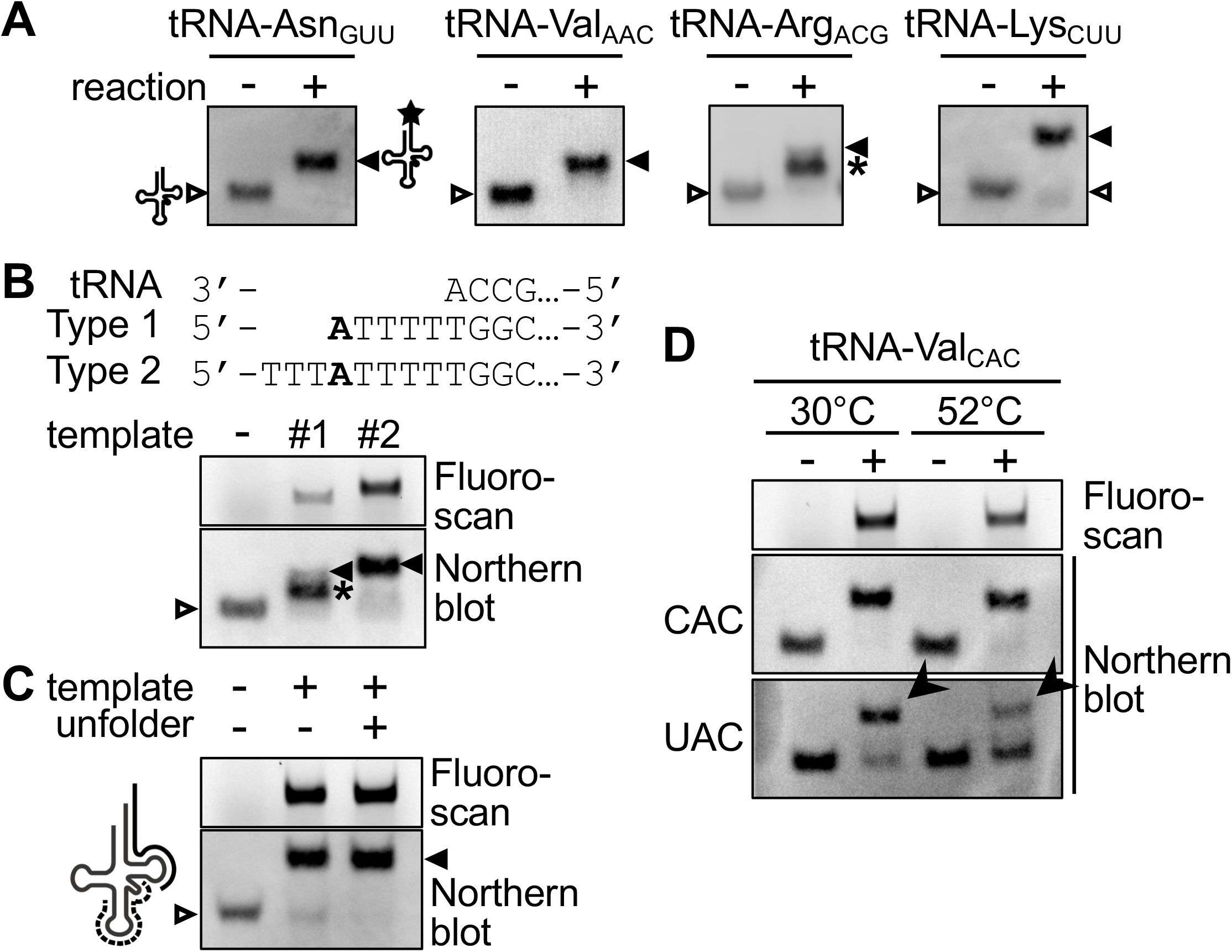
Optimization of OTTER reactions. (A) Northern blot analysis of OTTER reactions of tRNA-Asn_GUU_, tRNA-Val_AAC_, tRNA-Arg_ACG_, tRNA-Lys_CUU_ to elucidate labeling efficiency. OTTER reactions were performed in the absence (−) or presence (+) of corresponding Type 1 oligo DNA templates. Unextended tRNAs (open triangle), fully labeled species (closed triangle) and reaction intermediates (asterisk) were indicated. (*B*) OTTER reaction with internal labeling. The full sequence of each oligo DNA was shown in Table S1. OTTER reactions of tRNA-Arg_ACG_ with a Type1 template (#1), a Type 2 template (#2) or none (−) were separated in an 8% polyacrylamide gel, and were detected by fluorescence scanning (upper) and by Northern blotting (lower). (*C*) Effect of the unfolder oligo DNA on OTTER reactions. An unfolder oligo DNA (dashed line in the schematic drawing on the left; Table S1) was designed complementary to the anticodon stem-loop and variable loop region of tRNA-Lys_CUU_. OTTER reactions in the absence (−) or presence (+) of the unfolder were subjected to Northern blotting with a probe for tRNA-Lys_CUU_. (*D*) Effect of different annealing temperatures on OTTER reactions. OTTER reactions with a template oligo DNA for tRNA-Val_CAC_ were subjected to fluorescence scanning (top) and to Northern blotting with probes for tRNA-Val_CAC_ (middle) and for tRNA-Val_UAC_ (bottom). Bands marked by arrowheads correspond to tRNA-Val_UAC_ cross-reacted to the tRNA-Val_CAC_ template.

### tRNA quantification of yeast cells under several growth conditions

We applied OTTER to measure yeast tRNAs growing under several laboratory conditions. First, we measured and compared amounts of tRNAs in RNA samples prepared from the logarithmically growing phase. As shown in Fig. 3 and Table 1, tRNA amounts in the log-phase cells growing in a rich glucose medium, YPD, are in the range from 0.030 ± 0.002 pmol/μg RNA to 0.728 ± 0.044 pmol/μg RNA (24-fold difference). The least isoacceptor is tRNA-Leu_GAG_ encoded by a single gene while the most one is tRNA-Asp_GUC_ encoded by 16 synonymous genes. Total amount of tRNAs was 9.11 ± 0.56 pmol/μg RNA, which accounts for ~20% of total RNA in weight.

**Table 1.**
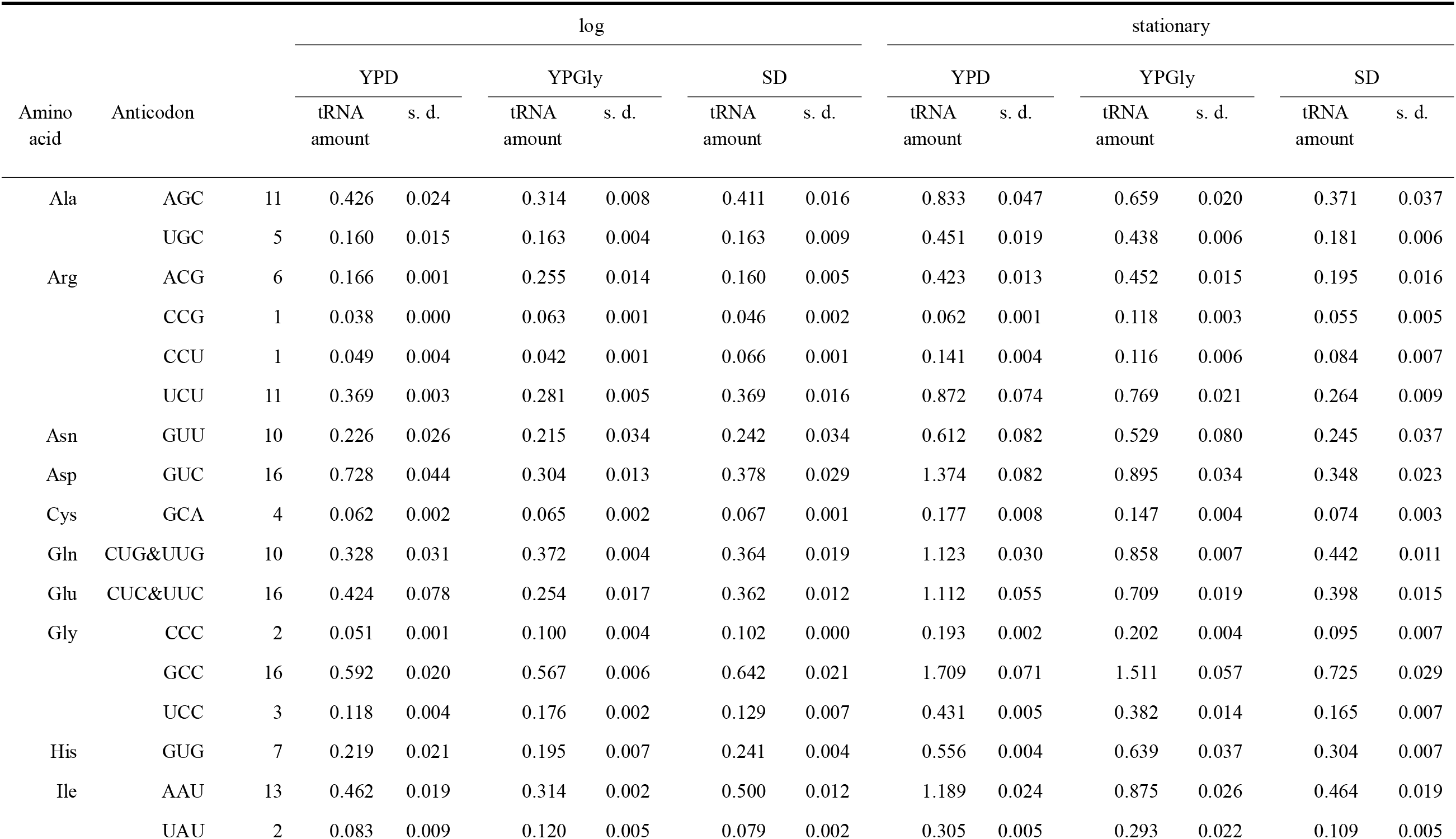
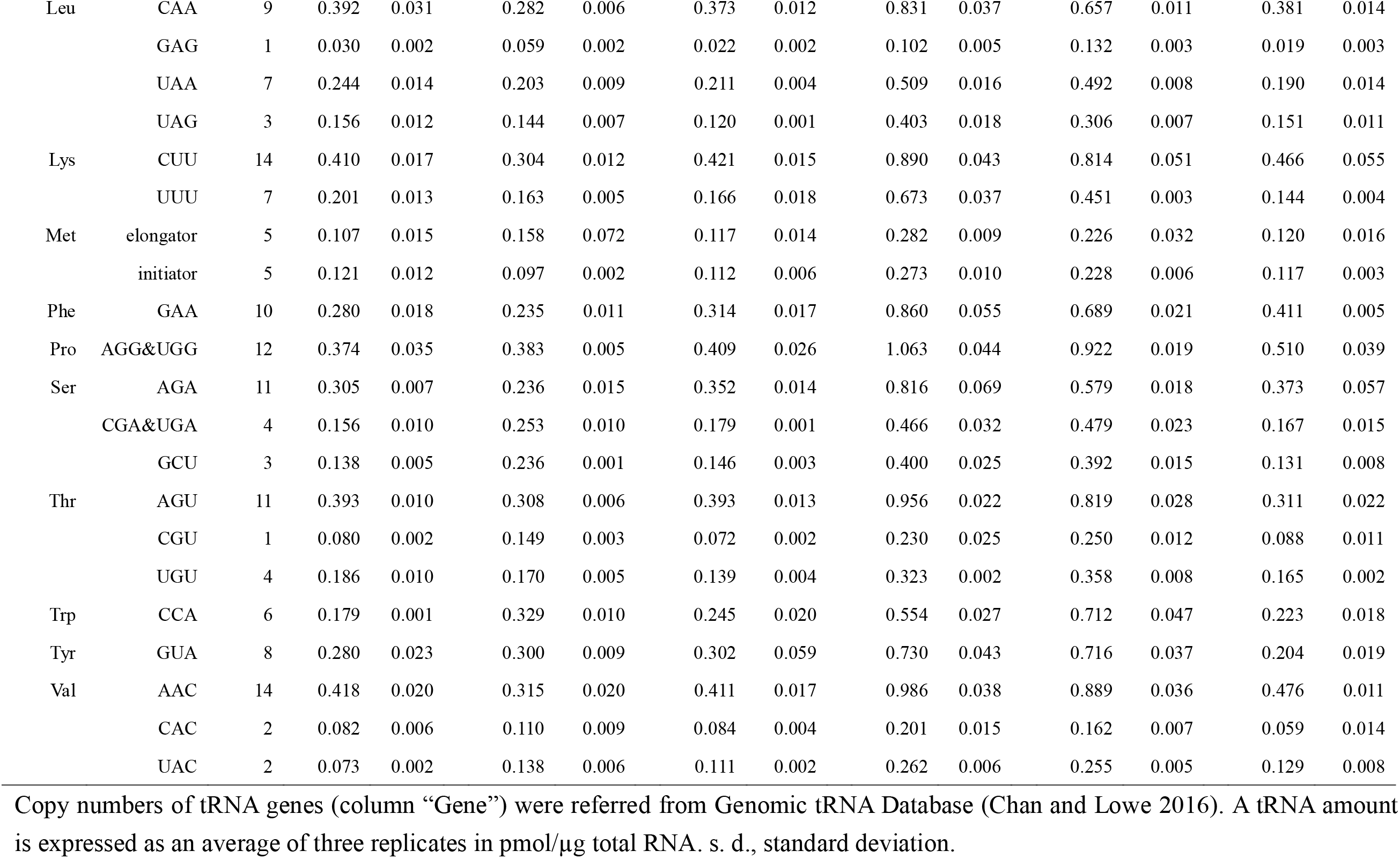
Summary of tRNA measurements by OTTER.

| Amino acid | Anticodon |  | log |  |  |  |  |  | stationary |  |  |  |  |  |
| --- | --- | --- | --- | --- | --- | --- | --- | --- | --- | --- | --- | --- | --- | --- |
|  |  |  | YPD |  | YPGly |  | SD |  | YPD |  | YPGly |  | SD |  |
|  |  |  | tRNA amount | s. d. | tRNA amount | s. d. | tRNA amount | s. d. | tRNA amount | s. d. | tRNA amount | s. d. | tRNA amount | s. d. |
| Ala | AGC | 11 | 0.426 | 0.024 | 0.314 | 0.008 | 0.411 | 0.016 | 0.833 | 0.047 | 0.659 | 0.020 | 0.371 | 0.037 |
|  | UGC | 5 | 0.160 | 0.015 | 0.163 | 0.004 | 0.163 | 0.009 | 0.451 | 0.019 | 0.438 | 0.006 | 0.181 | 0.006 |
| Arg | ACG | 6 | 0.166 | 0.001 | 0.255 | 0.014 | 0.160 | 0.005 | 0.423 | 0.013 | 0.452 | 0.015 | 0.195 | 0.016 |
|  | CCG | 1 | 0.038 | 0.000 | 0.063 | 0.001 | 0.046 | 0.002 | 0.062 | 0.001 | 0.118 | 0.003 | 0.055 | 0.005 |
|  | CCU | 1 | 0.049 | 0.004 | 0.042 | 0.001 | 0.066 | 0.001 | 0.141 | 0.004 | 0.116 | 0.006 | 0.084 | 0.007 |
|  | UCU | 11 | 0.369 | 0.003 | 0.281 | 0.005 | 0.369 | 0.016 | 0.872 | 0.074 | 0.769 | 0.021 | 0.264 | 0.009 |
| Asn | GUU | 10 | 0.226 | 0.026 | 0.215 | 0.034 | 0.242 | 0.034 | 0.612 | 0.082 | 0.529 | 0.080 | 0.245 | 0.037 |
| Asp | GUC | 16 | 0.728 | 0.044 | 0.304 | 0.013 | 0.378 | 0.029 | 1.374 | 0.082 | 0.895 | 0.034 | 0.348 | 0.023 |
| Cys | GCA | 4 | 0.062 | 0.002 | 0.065 | 0.002 | 0.067 | 0.001 | 0.177 | 0.008 | 0.147 | 0.004 | 0.074 | 0.003 |
| Gln | CUG&UUG | 10 | 0.328 | 0.031 | 0.372 | 0.004 | 0.364 | 0.019 | 1.123 | 0.030 | 0.858 | 0.007 | 0.442 | 0.011 |
| Glu | CUC&UUC | 16 | 0.424 | 0.078 | 0.254 | 0.017 | 0.362 | 0.012 | 1.112 | 0.055 | 0.709 | 0.019 | 0.398 | 0.015 |
| Gly | CCC | 2 | 0.051 | 0.001 | 0.100 | 0.004 | 0.102 | 0.000 | 0.193 | 0.002 | 0.202 | 0.004 | 0.095 | 0.007 |
|  | GCC | 16 | 0.592 | 0.020 | 0.567 | 0.006 | 0.642 | 0.021 | 1.709 | 0.071 | 1.511 | 0.057 | 0.725 | 0.029 |
|  | UCC | 3 | 0.118 | 0.004 | 0.176 | 0.002 | 0.129 | 0.007 | 0.431 | 0.005 | 0.382 | 0.014 | 0.165 | 0.007 |
| His | GUG | 7 | 0.219 | 0.021 | 0.195 | 0.007 | 0.241 | 0.004 | 0.556 | 0.004 | 0.639 | 0.037 | 0.304 | 0.007 |
| Ile | AAU | 13 | 0.462 | 0.019 | 0.314 | 0.002 | 0.500 | 0.012 | 1.189 | 0.024 | 0.875 | 0.026 | 0.464 | 0.019 |
|  | UAU | 2 | 0.083 | 0.009 | 0.120 | 0.005 | 0.079 | 0.002 | 0.305 | 0.005 | 0.293 | 0.022 | 0.109 | 0.005 |
| Leu | CAA | 9 | 0.392 | 0.031 | 0.282 | 0.006 | 0.373 | 0.012 | 0.831 | 0.037 | 0.657 | 0.011 | 0.381 | 0.014 |
|  | GAG | 1 | 0.030 | 0.002 | 0.059 | 0.002 | 0.022 | 0.002 | 0.102 | 0.005 | 0.132 | 0.003 | 0.019 | 0.003 |
|  | UAA | 7 | 0.244 | 0.014 | 0.203 | 0.009 | 0.211 | 0.004 | 0.509 | 0.016 | 0.492 | 0.008 | 0.190 | 0.014 |
|  | UAG | 3 | 0.156 | 0.012 | 0.144 | 0.007 | 0.120 | 0.001 | 0.403 | 0.018 | 0.306 | 0.007 | 0.151 | 0.011 |
| Lys | CUU | 14 | 0.410 | 0.017 | 0.304 | 0.012 | 0.421 | 0.015 | 0.890 | 0.043 | 0.814 | 0.051 | 0.466 | 0.055 |
|  | UUU | 7 | 0.201 | 0.013 | 0.163 | 0.005 | 0.166 | 0.018 | 0.673 | 0.037 | 0.451 | 0.003 | 0.144 | 0.004 |
| Met | elongator | 5 | 0.107 | 0.015 | 0.158 | 0.072 | 0.117 | 0.014 | 0.282 | 0.009 | 0.226 | 0.032 | 0.120 | 0.016 |
|  | initiator | 5 | 0.121 | 0.012 | 0.097 | 0.002 | 0.112 | 0.006 | 0.273 | 0.010 | 0.228 | 0.006 | 0.117 | 0.003 |
| Phe | GAA | 10 | 0.280 | 0.018 | 0.235 | 0.011 | 0.314 | 0.017 | 0.860 | 0.055 | 0.689 | 0.021 | 0.411 | 0.005 |
| Pro | AGG&UGG | 12 | 0.374 | 0.035 | 0.383 | 0.005 | 0.409 | 0.026 | 1.063 | 0.044 | 0.922 | 0.019 | 0.510 | 0.039 |
| Ser | AGA | 11 | 0.305 | 0.007 | 0.236 | 0.015 | 0.352 | 0.014 | 0.816 | 0.069 | 0.579 | 0.018 | 0.373 | 0.057 |
|  | CGA&UGA | 4 | 0.156 | 0.010 | 0.253 | 0.010 | 0.179 | 0.001 | 0.466 | 0.032 | 0.479 | 0.023 | 0.167 | 0.015 |
|  | GCU | 3 | 0.138 | 0.005 | 0.236 | 0.001 | 0.146 | 0.003 | 0.400 | 0.025 | 0.392 | 0.015 | 0.131 | 0.008 |
| Thr | AGU | 11 | 0.393 | 0.010 | 0.308 | 0.006 | 0.393 | 0.013 | 0.956 | 0.022 | 0.819 | 0.028 | 0.311 | 0.022 |
|  | CGU | 1 | 0.080 | 0.002 | 0.149 | 0.003 | 0.072 | 0.002 | 0.230 | 0.025 | 0.250 | 0.012 | 0.088 | 0.011 |
|  | UGU | 4 | 0.186 | 0.010 | 0.170 | 0.005 | 0.139 | 0.004 | 0.323 | 0.002 | 0.358 | 0.008 | 0.165 | 0.002 |
| Trp | CCA | 6 | 0.179 | 0.001 | 0.329 | 0.010 | 0.245 | 0.020 | 0.554 | 0.027 | 0.712 | 0.047 | 0.223 | 0.018 |
| Tyr | GUA | 8 | 0.280 | 0.023 | 0.300 | 0.009 | 0.302 | 0.059 | 0.730 | 0.043 | 0.716 | 0.037 | 0.204 | 0.019 |
| Val | AAC | 14 | 0.418 | 0.020 | 0.315 | 0.020 | 0.411 | 0.017 | 0.986 | 0.038 | 0.889 | 0.036 | 0.476 | 0.011 |
|  | CAC | 2 | 0.082 | 0.006 | 0.110 | 0.009 | 0.084 | 0.004 | 0.201 | 0.015 | 0.162 | 0.007 | 0.059 | 0.014 |
|  | UAC | 2 | 0.073 | 0.002 | 0.138 | 0.006 | 0.111 | 0.002 | 0.262 | 0.006 | 0.255 | 0.005 | 0.129 | 0.008 |

Copy numbers of tRNA genes (column “Gene”) were referred from Genomic tRNA Database (Chan and Lowe 2016). A tRNA amount is expressed as an average of three replicates in pmol/μg total RNA. s. d., standard deviation.

**Figure 3.**
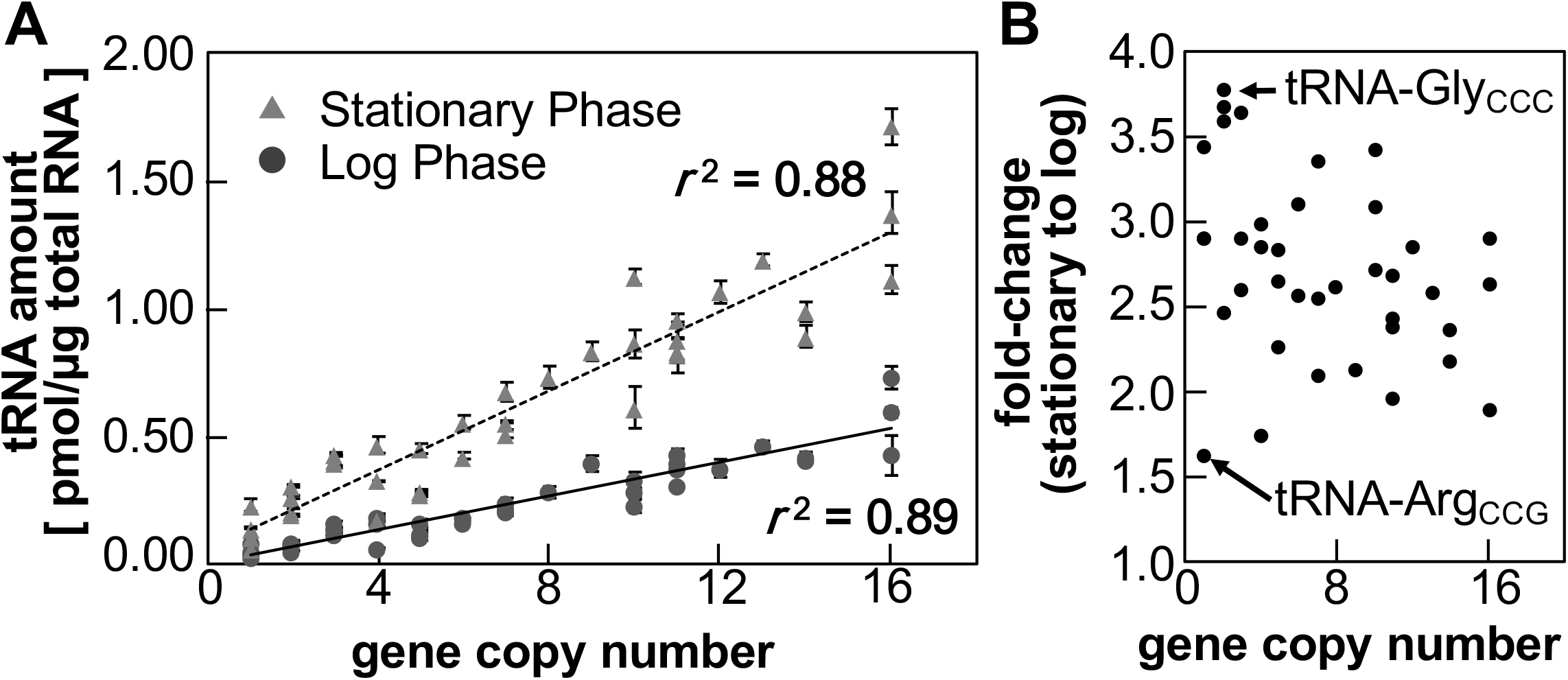
tRNA repertoires in yeast cells grown in the rich glucose medium. (*A*) The absolute amounts of tRNAs in RNA samples extracted from yeast cells growing to the log (circle) or stationary phase (triangle) were measured by OTTER. Each data set represents the average of 3 biological replicate. The error bar shows the standard deviation. tRNA gene copy numbers were referred from Genomic tRNA database (Chan and Lowe 2016). (*B*) Fold change in tRNA abundances in the stationary phase to the log phase. The tRNA species with the least change (tRNA-Arg_CCG_) and that with the highest change (tRNA-Gly_CCC_) were indicated by arrows.

Absolute amounts of tRNAs are far more closely related to the number of synonymous genes than we had expected; a correlation coefficient (*r*^2^) is 0.89 (Fig. 3A). On the other hand, tRNA amounts per gene vary between 0.016 pmol/μg RNA/gene for tRNA-Cys_GCA_ and 0.080 pmol/μg RNA/gene for tRNA-Thr_CGU_, and the average is 0.035 pmol/μg RNA/gene in the log phase yeast, meaning that there is about 5-fold difference in tRNA expression per gene.

Then, we compared tRNA amounts under different growth conditions. We measured amounts of tRNAs in the yeast cells growing in the rich medium with a non-fermentable carbon source, glycerol (YPGly), and those in the synthetic glucose medium (SD). We took RNA samples from cells both in the logarithmically growing phase and in the stationary phase to see the effect of growth phase progression. The results are summarized in Table 1. First, we looked closer at effects of the growth phase in the YPD culture. As shown in Fig. 3A, tRNA amounts per total RNA increased significantly in the stationary phase. Again, the tRNA amounts and the gene copy number are well-correlated (Fig. 3A; *r*^2^ = 0.88). However, changes in tRNA abundance from the log phase to the stationary phase vary from 1.6-fold (tRNA-Arg_CCG_) to 3.8-fold increase (tRNA-Gly_CCC_), an average of fold-changes is 2.7 (Fig. 3B, Table 1). This observation directly shows that amounts of tRNAs change according to cellular proliferative states and that the amount of each isoacceptor tRNA is somehow altered individually in the range of a few folds. Similar increase in tRNA amounts in the stationary phase was also observed when the cells were cultured in YPGly; 2.3-fold increase on average (from 1.4-fold in tRNA-Met_elongator_ to 3.3-fold in tRNA-His_GUG_). Interestingly, such increase was not obvious when the yeast cells were grown in the SD medium while amounts of tRNAs in the log-phase cells cultured in SD were comparable to those cultured in YPD and YPGly (Table 1).

Next, we investigated difference of tRNA amounts in log-phase cells cultured in the different media. As visualized in Fig. 4A, no extraordinary change was seen among these three medium conditions including total amounts of tRNAs (9.11, 8.37, and 9.02 pmol/μg RNA in YPD, YPGly, and SD, respectively). However, when investigated in detail, some characteristics of changes in tRNA repertoires were revealed. First, we compared total amounts of tRNAs for a certain amino acid. Yeast cells growing in YPGly and SD had about 40–50% of tRNA-Asp, which consists of a single isoacceptor tRNA-Asp_GUC_, if compared with those growing in YPD while the amounts of most of the tRNA species changed only in the rage of ± 15% (see Table 1, in detail). Similar, although milder, variation was seen in tRNA-Glu, consisting of tRNA-Glu_CUC_ and tRNA-Glu_UUC_ isoacceptors. On the other hand, tRNA-Trp, with a single tRNA-Trp_CCA_ isoacceptor, behaved oppositely; 1.8-fold more tRNA-Trp_CCA_ existed in the yeast cells growing in YPGly than those in YPD, and 1.4-fold more in SD. Even in the isoacceptor level, there are some interesting differences. For example, tRNA-Leu comprises one minor isoacceptor, tRNA-Leu_GAG_, and 3 major isoacceptors, tRNA-Leu_CAA_, tRNA-Leu_UAA_ and tRNA-Leu_UAG_ (Fig. 4B, left). When comparing the mounts in the YPD-grown cells to those in the YPGly-grown cells, the mounts of the three major isoacceptors decreased slightly in the latter cells (72–92% of those in the YPD-grown cells), like the total amount of tRNA-Leu (from 0.82 pmol/μg RNA to 0.69 pmol/μg RNA). On the other hand, the minor isoacceptor tRNA-Leu_GAG_ increased nearly 2-fold. Thus, the isoacceptor ratio of tRNA-Leu_GAG_ in the Leu-charged tRNAs increased from 3.6% in YPD to 8.6% in YPGly. Similar counter-acting behavior between major and minor isoacceptors was also observed in other tRNAs including tRNA-Thr (tRNA-Thr_AGU_ and tRNA-Thr_UGU_ vs tRNA-Thr_CGU_) and tRNA-Val (tRNA-Val_AAC_ vs tRNA-Val_UAC_) (Fig. 4B, center and right). And any of these cases allows about 2-fold increase of minor tRNA contribution to decoding of the particular amino acid under the respiratory conditions. The results indicate that tRNA repertoire are varied both in the amino acid isoform level and in the isoacceptor level according to carbon sources, and that the range of variation is about 40–200% of the tRNA amounts in the YPD-grown cells. This may be active regulation of tRNA repertoire in the yeast.

**Figure 4.**
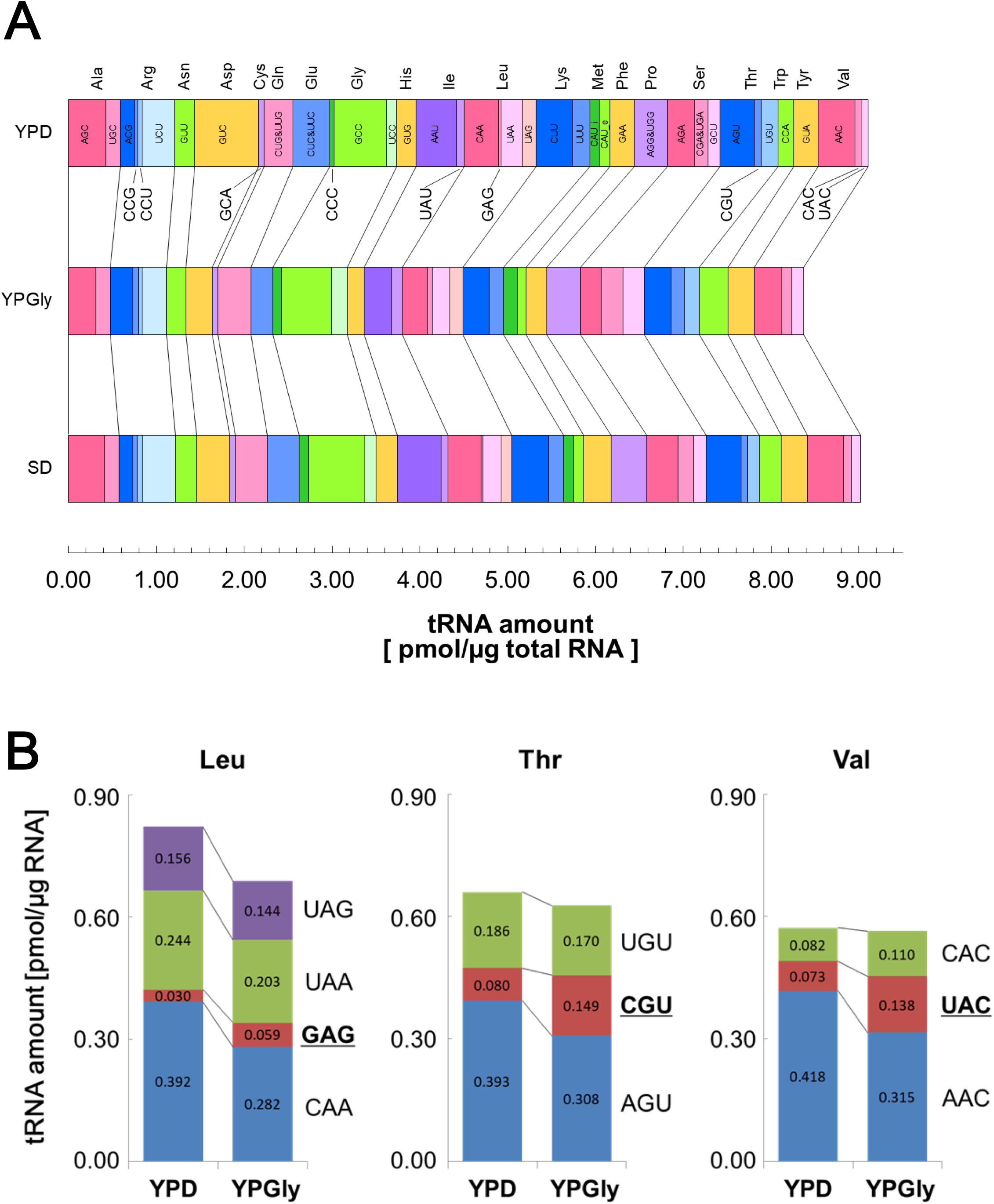
Comparison of tRNA amounts in the yeast cells growing in different media. (*A*) Comparison of tRNA mounts in the RNA samples prepared from log phase cells grown in YPD (top), YPGly (middle) and SD (bottom). The width of each bar represents the absolute amount of the indicated isoacceptor tRNA. (*B*) Alteration of isoacceptor contents grown in different carbos sources. tRNA amounts of isoacceptors for tRNA-Leu (left), tRNA-Thr (center), and tRNA-Val (right) in yeast cells grown either in YPD or in YPGly were shown as bar graphs.

### Comparison between OTTER and other quantification methods

Finally, we compared our tRNA quantitation results with other data obtained by different methods. First, we took two cases, RNA-Seq analysis by Dedon’s group (Pang et al. 2014) and microarray analysis by Pilpel’s group (Tuller et al. 2010), from high-throughput tRNA quantification studies. Essentially, there is no correlation between tRNA amounts determined by OTTER and RNA-Seq reads (Fig. 5A; *r*^2^ = 0.02), and moderate correlation is observed between our data and those from the microarray (Fig. 5B; *r*^2^ = 0.48). Since both OTTER and microarray rely on hybridization but not on reverse transcription, the big difference from the RNA-Seq analysis probably comes from unevenness of cDNA synthesis among different tRNA species caused by nucleotide modifications and/or tRNAs’ tight secondary structure. Indeed, the correlation efficient between the RNA-Seq data and the microarray data is also very low (*r*^2^ = 0.002). We did one more comparison of our data with the report by Ikemura (Ikemura 1982), where RNA samples prepared from ^32^P-labeled yeast were separated by 2D-urea PAGE and measured radioactivities of identified tRNA spots. Although this method only resolved 21 out of 42 total tRNA species (one of the 20 tRNA spots on the 2D gel was a mixture of tRNA-Ser_CGA_ and tRNA-Ser_UGA_), the quantification principle is quite simple, and has the least compromises and complexities in its experimental/data processing procedures among the methods analyzing here including ours. As shown in Fig. 5C, quantification by OTTER shows high correlation (*r*^2^ = 0.80) to the results of the 2D-PAGE method. We also compare tRNA quantitation by the 2D-Gel with the other three methods using data of 17 tRNA species common to all the four methods (including the sum of tRNA-Ser_CGA_ and tRNA-Ser_UGA_). Correlation coefficients to OTTER, RNA-Seq, and microarray are 0.75, 0.05, and 0.34, respectively, and OTTER shows the best correlation to the 2D-PAGE data among the three. Thus, these results indicate that OTTER is the most suitable method to compare amounts of different tRNA species.

**Figure 5.**
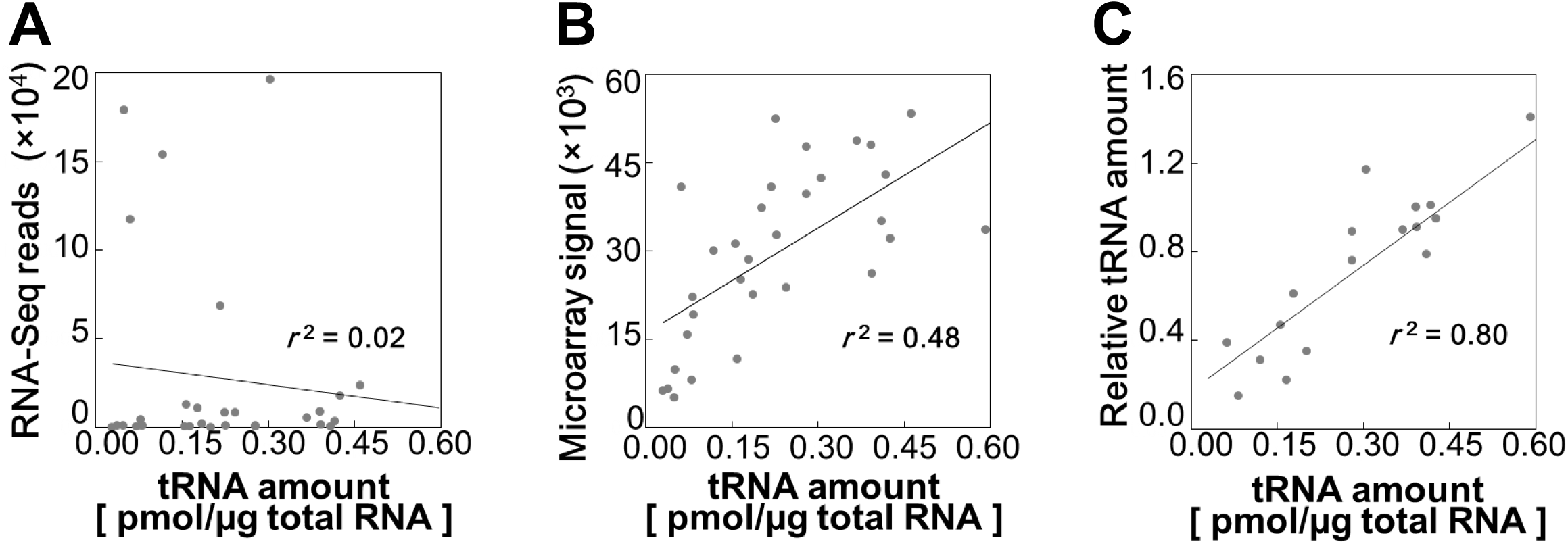
Comparison between OTTER and other tRNA quantification methods. The absolute amounts of isoacceptor tRNAs in the RNA sample from logarithmically growing cells in the glucose medium were measured by OTTER and compared with the relative tRNA levels in the previously reported tRNA-Seq data (Pang et al. 2014) (*A*), the microarray data (Tuller et al 2010) (*B*), and the 2D-PAGE data (Ikemura 1982) (*C*).

## DISCUSSION

### A new method for tRNA quantification, OTTER; its pro and con

Here, we introduced a new quantification method for tRNA, OTTER (Fig. 1). It directly fluorescence-labels a specific tRNA species according to sequence diversity of its 3’ region. Because our OTTER includes the assessment procedure of signal production efficiency and cross-reactivity through Northern analysis, this method circumvents the problem of variation in signal generation among different tRNA species/sample conditions. Indeed, as shown in Fig. 5, quantification data of OTTER showed good agreement to those obtained by ^32^P *in vivo*-labeling and quantitative 2D-gel analysis (*r*^2^ = 0.80) (Ikemura 1982), the most straightforward method for tRNA quantification. On the other hand, our data poorly correlated with the simple RNA-Seq results (*r*^2^ = 0.02) (Pang et al. 2014), partly because of uneven reverse transcription caused by variable nucleotide modifications and/or secondary structural stability of individual tRNAs. The moderate correlation between OTTER’s data and tRNA microarray’s data (*r*^2^ = 0.48) (Tuller et al. 2010) also supports this notion. On the other hand, what causes the difference between OTTER and the microarray? Although probes in microarray analysis are carefully designed, their hybridization efficiency and specificity cannot be evaluated one-by-one on the array. As shown in Fig. 2, DNA probes for OTTER hybridized to their cognate tRNAs with different efficiency. Certain tRNA pairs with similar 3’ sequences cannot be resolved (see below in detail) in OTTER despite the fact that such pairs harbor 2 nt mismatches in the middle part of their region hybridized with the OTTER probes for the non-cognate tRNAs. We speculate that microarray may also suffer from similar problems. Thus, OTTER is superior to RNA-Seq and microarray from the view of precision and by the fact that absolute amounts of tRNA species can be directly handled.

In our experience, efficiency of fluorescence labeling of a certain isoacceptor tRNA did not change so much among RNA samples from yeast cells grown under different physiological conditions. Therefore, a correction coefficient determined by a standard sample, such as an RNA sample from logarithmically growing yeast in the YPD medium, can be applied to other measurements when the highest precision is not required for the particular estimation of tRNA repertoire, which allows omission of Northern blotting in each sample as far as cross-reactivity is not the problem. In addition, OTTER can be applied to measure only a subset of tRNA species, so that it is also a versatile method for tRNA quantification in various analytical scenes.

There are several drawbacks in OTTER: one is limitation of tRNA resolution because our method fully relies on the diversity of 3’ region of tRNAs. Indeed, we cannot resolve yeast tRNA-Ser_CGA_ and tRNA-Ser_UGA_ in principle. We could not resolve Gln_CUG_/tRNA-Gln_UUG_, tRNA-Glu_CUC_/tRNA-Glu_UUC_, and tRNA-Pro_AGG_/tRNA-Pro_UGG_ pairs in practice, despite the fact that all of these tRNA pairs have some difference in their sequences in this region. Thus, RNA-Seq and even microarray are superior to our method in tRNA resolution. Another drawback comes from the fact that no amplification step is integrated into OTTER although it is required for precise determination of absolute amounts of tRNAs. Thus, we need some amount of RNA samples for quantification of a full-set of tRNAs.

### Quantification results of yeast tRNAs

OTTER revealed that the tRNA species in the log phase yeast cells exist in the range of 0.030–0.73 pmol/μg RNA, meaning that yeast cells have ~9 pmol/μg RNA of tRNAs in total (~20% of total RNA in weight). Since one yeast cell is estimated to have ~0.5 pg RNA (Boehlke and Friesen 1975), molecular numbers of individual tRNA species are calculated as 8,000–220,000 molecules/cell. Thus, supposing that most of the yeast tRNAs are localized in the cytosol (Sarkar and Hopper, 1998; Grosshans, Hurt and Simos, 2000), their cytosolic concentrations can be estimated as 0.55–13 μM from the average haploid cell volume and the average cytoplasm/cell ratio (Jorgensen et al. 2002; Yamaguchi et al. 2011). These *in vivo* parameters estimated from our data can be compared with numbers related to translation in yeast that were acquired previously through various biochemical and cell biological studies. For example, a yeast cell contains ~190,000 ribosomes (von der Haar 2008) and more than 500,000 molecules of eEF1A (Norbeck and Blomberg 1997). This means that, for the least tRNA species such as tRNA-Leu_GAG_, tRNA/ribosome ratio is less than 1/20 while molecular numbers of major tRNAs, such as tRNA-Asp_GUC_, are comparable to that of ribosomes. Therefore, these *in vivo* numbers support the notion that there is competition between ribosomes for acquiring minor tRNAs to their A site while such competition is quite mild, if not at all, in the case of major tRNAs. Each tRNA species must be charged with its cognate amino acid by aminoacyl-tRNA synthetase (ARS). Affinities of yeast ARSs to the cognate tRNAs were biochemically determined as a range from a few tens to a few hundreds nM (for example, 12 nM, 130 nM and 220 nM for AspRS, TyrRS and SerRS, respectively) (Erinai et al. 1991; Soma and Himeno 1998). The estimated range of tRNA concentrations from our data is higher than these numbers, and it is consistent to the fact that most of the tRNA species are almost fully aminoacylated in the normal growth conditions (Zaborske et al. 2009).

The absolute amounts of tRNAs we measured in yeast cells showed good correlation with the number of synonymous genes on the yeast genome. Thus, it is feasible that an amount of a certain tRNA can be estimated roughly from its gene copy number as adopted by many researchers. On the other hand, we noticed that tRNA amounts change in some ranges according to difference in physiological conditions. First, tRNA amounts per total RNA increase 2–3-fold when entered into the stationary phase. This was seen both in YPD and YPGly but not in SD. Although the exact substance(s) induces the phenomena is unknown, we speculate that shortage of some nutrients, such as nitrogen sources but not carbon sources, causes this increase. This much of increase in tRNA amounts per total RNA does not seem to come from upregulation of tRNA production because both RNA polymerases I and III are repressed in the stationary phase. However, repression of rRNA production precedes that of RNA polymerase III (Clarke et al. 1996), and this may contribute to the relative accumulation of tRNAs. In addition, ribosomal RNAs may be degraded under these conditions more efficiently than tRNAs partly via autophagy (Iwama and Ohsumi 2019). Because autophagy in the stationary phase seems to degrade bulk of the cytosol, tRNAs might somehow avoid autophagy. Or ribosomes are selectively degraded by so-called ribophagy in the stationary phase (Wyant et al. 2018).

Another important point is that individual tRNA species respond differently to the environmental changes. First, as shown in Fig. 3 and Table 1, increase in tRNA amounts during the progression of the growth phase vary from 1.6-fold to 3.8-fold in YPD and from 1.4-fold to 2.3-fold in YPGly, indicating that individual tRNA species seem to be differently regulated in the range of a few folds. It is to be noted that major and minor isoacceptor tRNAs of tRNA-Leu, tRNA-Thr *etc.* behave oppositely between YPD and YPGly (Fig. 4), implying that major/minor isoacceptor ratio is regulated according to minimize change of the total amount of the isoacceptor tRNAs for one amino acid. This suggests that isoacceptors for a certain amino acid are regulated individually but the regulation is coordinated. Although we still do not know whether such regulation is done in the transcriptional level or in the post-transcriptional (degradation) level, tRNA expression may be under more complex and sophisticated regulation than we expected.

## MATERIALS AND METHODS

### Yeast cell culture

*S. cerevisiae* BY418 [*MAT*α *ade2*Δ*::hisG his3*Δ*200 leu2*Δ*1 lys2*Δ*202 met15*Δ*0 trp1*Δ*63 ura3-52*] was grown at 30°C in liquid media YPD [1.0%(w/v) yeast extract, 2.0%(w/v) polypeptone, and 2.0%(w/v) D-glucose], YPGly [1.0%(w/v) yeast extract, 2.0%(w/v) polypeptone, and 2.0%(w/v) glycerol), or SD [0.67%(w/v) yeast nitrogen base without amino acids and 2.0%(w/v) D-glucose] with 20 μg/ml appropriate amino acid and nucleobase supplements. Cell growth was monitored by measuring OD_660_ using a Miniphoto518R spectrophotometer (Taitec, Koshigaya, Japan). The cells were harvested at the log phase (OD_660_ ≈ 0.3) or at the stationary phase by centrifugation at 1,610 ×*g* for 3 min. The cell pellets were quickly frozen in liquid nitrogen.

### Total RNA isolation

Total RNA extraction from yeast cells was performed with SDS under acidic conditions (Hayashi et al. 2019). Frozen yeast cells were resuspended in AES Buffer [50 mM sodium acetate, pH 5.2, 10 mM EDTA, 1.0%(w/v) SDS], and was extracted with acidic phenol chloroform [phenol:chloroform = 5:1, pH 4.5] at 65°C for 10 min with occasional mixing, and the water phase was separated by cooling in ice water and subsequent centrifugation. After extraction of the aqueous phase with phenol chloroform [phenol:chloroform = 1:1 with 0. 050%(w/v) 8-oxyquinoline] followed by chloroform extraction, RNAs were precipitated with 2-propanol, and the final pellet was dissolved in TE [10 mM Tris-HCl, pH 7.5, and 1.0 mM EDTA].

### Fluorescence labeling of tRNA

Template oligo DNAs shown in Table S1 were synthesized by Eurofins Genomics (Tokyo, Japan). Two μg of yeast total RNA was incubated with an appropriate oligo DNA template (12.5 pmol) in 8 μl of Assay Buffer [10 mM Tris-HCl, pH 8.0, 50 mM NaCl, 0.50 mM EDTA, 1.0 mM DTT] at 94°C for 3 min. The mixture was gradually cooled to 30°C and 52°C in 90 min and 59 min, respectively, and then quickly chilled at 4°C. After adding final concentration of 10 μM TMR-dUTP (Roche Diagnostics, Basel, Switzerland), 250 μM dATP, 10 mM MgCl_2_, and 2.0 units of Klenow Fragment (3’-5’ exo^−^)(New England Biolabs, Ipswich, Massachusetts, USA), total 10 μl of the mixture was incubate at 37°C for 90 min. The labeling reaction was stopped by mixing with 0.40 μl of 0.50 M EDTA. Combinations of template oligo DNA, annealing temperature, and data processing for individual tRNA species were summarized in Table S2.

### Quantification of fluorescence-labeled tRNAs

RNAs in the assay mixtures were separated by electrophoresis on an 8.0%(w/v) polyacrylamide gel containing 7.0 M urea in TBE buffer. Fluorescence of the labeled tRNAs on the gel was detected by a laser scanner, Typhoon FLA-7000 (GE Healthcare, Chicago, Illinois, USA), and the data were processed by a software, Image Quant TL (GE Healthcare). The absolute amount of each tRNA species was determined by using a known concentration of TMR-labeled oligo DNA (Sigma Genosys, St. Louis, Missouri, USA) as a standard substance.

### Northern blotting

For accurate measurement, labeling efficiency and cross-reactivity to homologous tRNAs of each target tRNA species were assessed by Northern blotting. RNAs in the urea-polyacrylamide gel were transferred to a charged nylon membrane, Hybond-N^+^ (GE Healthcare). A particular tRNA was hybridized with a corresponding antisense oligo DNA probe shown in Table S1 labeled with digoxigenin (DIG) using DIG Oligonucleotide Tailing Kit (Roche Diagnostics) in Hybridization Solution [0.50 M Na_2_HPO_4_, 0.34%(v/v) H_3_PO_4_, 7.0%(w/v) SDS, 1.0 mM EDTA, pH 7.0] at 37°C. After stringent wash in 4×SSC at 37°C, the hybridized DIG probe was decorated with alkaline phosphatase-conjugated anti-DIG antibodies (Roche Diagnostics), and was detected with CDP-Star chemiluminescence. The signal was captured and analyzed by a cooled CCD camera system, Ez-Capture and imaging analysis software, CS Analyzer 3 (ATTO, Tokyo, Japan).

## ACKNOWLEDGMENTS

We thank to our current and previous lab members for their support and discussion. This work is supported by KAKENHI for Scientific Research (C) Grant Numbers JP17K07289, JP17KT0113, and JP20K06490 from Japan Society for the Promotion of Science, and by KAKENHI for Scientific Research on Innovative Areas Grant Number JP17H05672 and JP20H05338 from Ministry of Education, Culture, Sports, Science and Technology Japan.

